# Diet alters micronutrient pathways in the gut and placenta that regulate fetal growth and development in pregnant mice

**DOI:** 10.1101/767012

**Authors:** Elia Palladino, Tim Van Mieghem, Kristin L. Connor

## Abstract

Maternal malnutrition and micronutrient deficiencies can alter fetal development. However, the mechanisms underlying these relationships are poorly understood. We used a systems-physiology approach to investigate diet-induced effects on maternal gut microbes and folate/inositol transport in the maternal/fetal gut and placenta. Female mice were fed a control diet (CON) diet, undernourished (UN, restricted by 30% of CON intake) or a high fat diet (HF, 60% kcals fat) during pregnancy to model normal pregnancy, fetal growth restriction, or maternal metabolic dysfunction, respectively. At gestational day 18.5 we assessed circulating folate levels by microbiological assay, relative abundance of gut lactobacilli by G3PhyloChip™, and folate/inositol transporters in placenta and maternal/fetal gut by qPCR/immunohistochemistry. UN and HF-fed mothers had lower plasma folate concentrations vs. CON. Relative abundance of three lactobacilli taxa were higher in HF vs. UN and CON. HF-fed mothers had higher gut proton coupled folate transporter (*Pcft*) and reduced folate carrier 1 (*Rfc1)*, and lower sodium myo-inositol co-transporter 2 (*Smit2)*, mRNA expression vs. UN and CON. HF placentae had increased folate receptor beta (*Fr*β) expression vs. UN. mRNA expression of *Pcft, folate receptor alpha (Frα)* and *Smit2* was higher in gut of HF fetuses vs. UN and CON. Transporter protein expression was not different between groups. Maternal malnutrition alters abundance of select gut microbes and folate/inositol transporters, which may influence maternal micronutrient status and delivery to the fetus, impacting pregnancy/fetal outcomes.

## Introduction

Folate and myo-inositol are two critical micronutrients for fetal central nervous system development. Folate plays an important role in spinal closure and folate deficiency increases the risk of neural tube defects [1–3]. Inositol, a pseudovitamin of the vitamin B family, is necessary for neurulation and essential for brain development [4–6]. Suboptimal maternal nutrition, which is seen in both underweight and obese pregnant women, is associated with reduced serum folate concentrations, even if the diet is sufficient in folate and inositol is an important insulin sensitizer which has been well documented in diabetic patients [7–12]. Despite the importance of folate and inositol, the extent to which malnutrition in pregnancy impacts production and transfer of these micronutrients from mother to fetus is unclear.

Folate is a naturally occurring essential B vitamin that is required for DNA synthesis and as such, is important during periods of rapid tissue growth, such as embryogenesis and fetoplacental development [13–15]. Folic acid is a synthetic form of folate found in dietary supplements [13]. Folate transfer to the fetus is mediated by the placental transporters folate receptor alpha and beta (FRα/β), proton coupled folate transporter (PCFT) and reduced folate carrier 1 (RFC1) [16,17]. FRα is the main folate transporter responsible for maternal-to-fetal transfer across the placenta (in both mice and humans) and works together with PCFT and RFC1 to control the uptake and release of folate [18–20]. PCFT is a high-affinity folate transporter and RFC1 mediates folate transport to the fetus via bi-directional transport [16,19,21–23]. Inositol is transferred to the fetus by placental transporters sodium (Na+) myo-inositol co-transporter 1 and 2 (SMIT1/2) and tonicity-responsive enhancer binding protein (TONEBP) [24,25]. SMIT1 is a high affinity transporter that is highly expressed prenatally in the fetal central nervous system and placenta [26]. Similarly, TONEBP is a main signal transcription factor of SMIT1 and the absence of TONEBP results in reduced *Smit1* transcription [25]. The SI also acts as a main regulator of folate/inositol uptake and transport via PCFT, RFC1, SMIT2, TONEBP. PCFT and RFC1 are expressed in the duodenum and jejunum, which are the preferred sites of folate absorption, and PCFT is responsible for the majority of folate intestinal absorption [16,27–29]. Moreover, SMIT2 is highly expressed in the intestinal epithelium of the gut, playing a greater role than SMIT1 in mediating gut inositol uptake/transport, opposite to their roles in the placenta [30]. Maternal gut folate and inositol absorption therefore is likely critical to the early supply of these micronutrients to the developing embryo/fetus, and throughout gestation [16,28,29,31]. To our knowledge, the presence of these transporters has not been investigated in the fetal gut. As such, their roles and importance to the fetus remain unclear.

In addition to dietary intake of nutrients, specific gut microbes play key roles in nutrient metabolism and vitamin production [32,33]. *Lactobacillus* and *Bifidobacterium* produce folate and increase production of these vitamins when paired with prebiotic fibres, therefore potentially compensating for the lack of dietary intake of folate in the host [27,32,34–40]. The small intestine (SI), particularly the duodenum, jejunum, and colon, are sites for bacterial folate production and absorption [27,41]. Although there is limited information on bacterial production of inositol, it is absorbed in the SI [42]. Once absorbed from the maternal gut into the blood, these micronutrients are then transferred transplacentally to the fetus [15,18,43,44].

Whilst maternal malnutrition, underweight, and overweight/obesity have been associated with reduced folate status and adverse pregnancy outcomes, the mechanisms through which these conditions lead to poor fetal development are not fully understood [1,2,7,8,10,11]. We have previously shown that maternal UN and HF diets in mice, used to recapitulate common pregnancy phenotypes of undernutrition/underweight and overweight/obesity (namely fetal growth restriction or maternal metabolic dysfunction, respectively), change the maternal and fetal gut barriers, the maternal gut microbiome and placental development and function [45–48]. These findings suggest that a systems physiology approach, considering the entire nutritional pipeline (maternal gut, placenta, fetal gut) is required to understand the mechanisms governing fetal development and growth. The present study therefore aimed to investigate how nutritional challenges impact circulating folate levels and folate/inositol pathways in the placenta and maternal and fetal guts. We hypothesised that maternal undernutrition and high fat diet would decrease circulating folate levels and result in pregnancy adaptations including increased abundance of maternal gut Lactobacillus and altered expression of genes and proteins in the maternal gut and placenta important for to folate/inositol production and transport. We also hypothesised that these adaptations would be associated with increased expression of folate and inositol pathways in the fetal gut.

## Materials and Methods

### Animal model and diet

All experiments were approved by the Animal Ethics Committee at Mount Sinai Hospital, Toronto, Canada (AUP-0091a-H). C57BL/6J female mice (Jackson Laboratories, Bar Harbour, ME, USA) were randomised to three nutritional groups: control (CON), undernourished (UN) and high fat (HF), designed to model changes seen in maternal and fetal physiology in humans with undernutrition/underweight and overweight/obesity[45–48]. Female mice were fed a control diet (23.4% saturated fat by weight; Dustless Precision Pellets S0173, BioServe, Frenchtown, NJ, USA) *ad libitum* before and throughout pregnancy (n=7), *or* mice were moderately undernourished restricting food by 30% of control intake from gestational day (GD) 5.5-17.5 of pregnancy (n=7) *or* mice were fed a 60% high fat diet (60% kcals as fat, 37.1% saturated fat by weight; D12492, Research Diets, New Brunswick, NJ, USA) from 8 weeks before mating and during pregnancy (n=8) [46]. Of note, there were differences in macronutrient and micronutrient content between diets (Table 1), as is also seen in the clinical human setting. Fathers (Jackson Laboratories, Bar Harbour, ME, USA) were fed a control diet *ad* libitum and were mated with females at approximately 10 weeks of age.

**Table 1.**
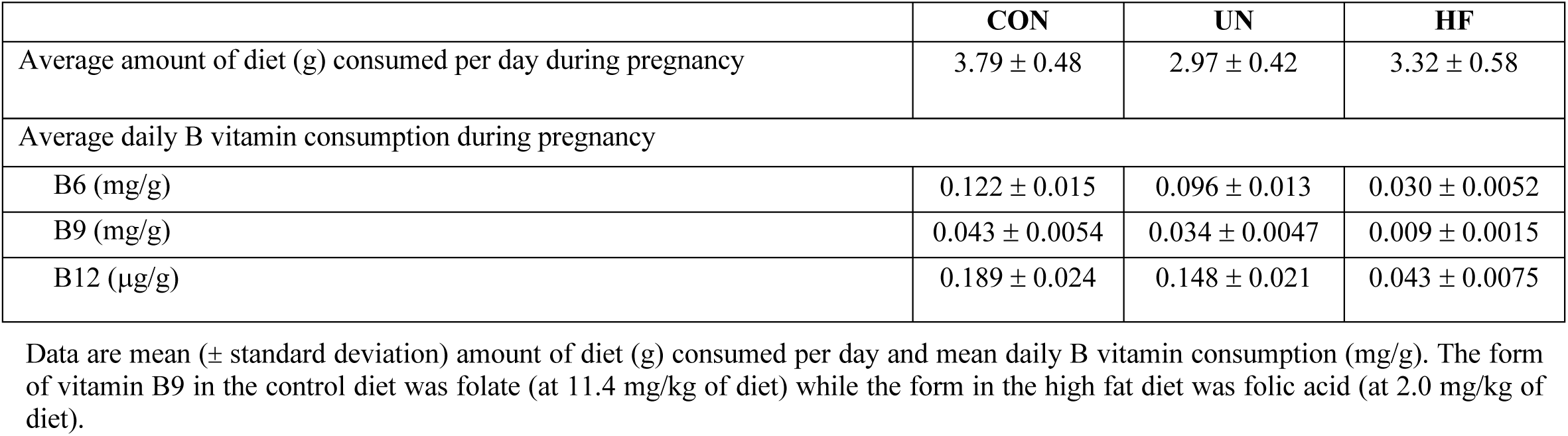
Composition of CON, UN and HF diets.

### RNA extraction and expression in placenta and gut

At GD18.5, maternal, placental, and fetal tissues were collected and snap frozen in liquid nitrogen and stored at −80 until analysed. RNA was extracted from whole jejunum homogenates from each mother (CON n=7, UN n=7, HF n=8), from whole placental homogenates from one male and one female placenta from each litter, and from whole fetal gut homogenates from one male and one female fetus from each litter (females: CON n=7, UN n=4, HF n=8 HF; males: CON n=6, UN n=6, HF n=8) using the Qiagen RNeasy Plus Mini Kit (Qiagen, Toronto ON). RNA was quantified by spectrophotometry (DeNovix, Wilmington, DE, USA) and 1 μg of RNA was reverse transcribed by using 5X iScript Reverse Transcription Supermix (Bio-Rad, Mississauga, ON, Canada) according to the manufacturer’s instructions.

Maternal jejunal SI mRNA expression of *Pcft and Rfc1*(folate transporters), *Smit2* and *TonEBP* (inositol transporters), and placental and fetal gut expression of *Frα, Frβ, Pcft, Rfc1, Smit1/2* and *TonEBP* were measured by real time qPCR using SYBR Green I (Bio-Rad, Mississauga ON) and 5 μM of forward and reverse primer mix (Bio-Rad CFX384, Mississauga ON, Canada). Primer sequences for genes of interest and three stably expressed reference genes were designed using Primer BLAST software (NCBI) or from the literature (Online Resource Table 1). A standard curve was run for each gene. Standards, samples, and controls were run in triplicate under the following cycling conditions: 95°C for 30 seconds, 60°C for 20 seconds, 39 successive cycles at 95°C for 5 seconds and 60° C for 20 seconds, and melt curve at 65°C with 0.5°C increments for 5 seconds/cycle. Target gene expression was normalised to the geometric mean of the following reference genes: Beta-actin *(Actb)*, Glyceraldehyde 3-phosphate dehydrogenase *(Gapdh)* and Hypoxanthine Phosphoribosyltransferase 1 (*Hprt1)* (placental samples); *Actb*, Tyrosine 3-Monooxygenase/Tryptophan 5-Monooxygenase Activation Protein Zeta (*Ywhaz)* and TATA Box Binding Protein *(TBP)* (maternal SI and fetal gut samples). Data were analysed using the Pfaffl method [49].

### Localisation of FRβ, PCFT and SMIT2 protein

To establish whether diet-induced pregnancy phenotypes were related to changes in the distribution of select micronutrient receptors and transporters in the maternal SI (intestinal mucosa and epithelial cells) and placenta (decidua, junctional and labyrinth zones), immunohistochemical (IHC) analyses were conducted. Placentae and SI samples were fixed in paraformaldehyde, paraffin embedded, and sections cut at 5-μm. To visualise FRβ immunoreactive distribution, placental midsections from 1 male and 1 female placenta per litter were stained using a rabbit polyclonal anti-FOLR2 antibody (Abcam, ab228643, Cambridge, United Kingdom) at a 1:50 dilution. To visualise PCFT and SMIT2 immunoreactive distribution, SI sections were stained with rabbit polyclonal anti-HCP1/PCFT antibody (Abcam, ab25134, Cambridge, United Kingdom) and rabbit polyclonal anti-SLC5A11/SMIT2 antibody (LifeSpan BioSciences, LS-C403570-120, Seattle, WA, USA). Slides were deparaffinised in 100% xylene, rehydrated in decreasing concentrations of ethanol, and quenched using 3.0% hydrogen peroxide in methanol at room temperature. Antigen retrieval was performed by microwave treatment of 10mM solution of citric acid and sodium citrate (pH 6.0). Slides were washed with 1X phosphate-buffered saline (PBS) and incubated in a humidified chamber for 1 hour at room temperature with protein-blocking solution (Agilent Dako, Santa Clara, ON, Canada) to block nonspecific sites. Slides were incubated overnight at 4°C with the specific primary antibody. A negative control was included in each run for each primary antibody (primary antibody omitted in replace of antibody diluent). Slides were incubated with 1:200 goat biotinylated anti-rabbit antibody (BA-1000, Vector Labs, Burlingame, CA, USA) in a humidified chamber for 1 hour at room temperature, followed by incubation with streptavidin-horse radish peroxidase (HRP, 1:2000 dilution in 1X PBS, Invitrogen, Carlsbad, CA, USA) for 1 hour at room temperature. To visualise HRP enzymatic activity, the reaction was initiated by 3,3’ diaminobenzidine (DAB; Vectastain Elite ABC HRP Kit, Vector Laboratories, Brockville, ON, Canada) for 3 minutes and 30 seconds. Slides were counterstained with Gill’s #1 haematoxylin (Sigma-Aldrich, Oakville, ON, Canada).

To evaluate the localisation and expression of immunoreactive (ir) FRβ, PCFT and SMIT2 stained sections, sections were semi-quantitatively analysed as previously described [50]. Images were taken on the EVOS FL Auto 2 (Thermo Fisher Scientific, Waltham, MA, USA) at 20X magnification by one researcher blinded to the animal IDs and groups. Entire placentae and SI sections were scanned to note staining patterns. Placental images were taken in the decidua, junctional and labyrinth layers and chorionic plate. Across all samples using the automate function (EVOS FL Auto 2, Thermo Fished Scientific, Waltham, MA, USA), six images were randomly captured within each layer of the placenta. Using the same function, six images were randomly captured along the length of each maternal small intestine section. Intensity of staining was scored based on the scoring system described previously by a single observer blinded to the experimental groups [50]. Staining intensity was classified on a 5-point scale: 0 = absent, 1 = weak, 2 = moderate, 3 = strong and 4 = very strong [50]. For all sections, average staining intensity across the six images was calculated to determine the mean intensity for each animal. Semi-quantitative staining intensity data are presented as mean intensity staining for the specified protein and diet group calculated from the average staining intensity across all six images.

### 16S rRNA gene sequencing and analysis

DNA was extracted from maternal caecal contents at GD18.5, as previously described.^[45]^ Diversity and relative abundance of maternal gut microbes were measured using the G3 PhyloChip™[46].

### Circulating folate measures

To determine maternal folate status, circulating folate levels were measured using the *Lactobacillus casei* microbiological assay on maternal plasma samples, as previously described [51,52].

### Statistical analysis

Analyses were performed using JMP 14 (SAS Institute Inc., Cary, NC, USA). Outcome measures were analysed for normality and equal variances (Levene’s test). Data that were normal and had equal variances were analysed by parametric tests. If data were non-normal, a log transformation was applied to normalise. Data that could not be normalised were analysed by non-parametric tests. Differences between dietary groups for outcome measures were compared by ANOVA (Tukey’s *post hoc*), or Welch ANOVA (Games-Howell *post hoc*), or Kruskal-Wallis (Steel-Dwass *post hoc*). Relationships between variables were determined by Pearson or Spearman correlation analysis. P = <0.05 was deemed statistically significant. Data are presented as means ± standard deviation (SD) or median interquartile range (IQR), with 95% confidence interval diamonds in figures.

## Results

### Malnutrition is associated with reduced maternal plasma folate levels and fetal weight

Previously we reported that maternal malnutrition altered maternal and fetoplacental weights [48]. In this study, maternal plasma folate levels were lower in both UN and HF fed mothers compared to CON mothers (p<0.0001, Fig. 1a), with HF fed females having the lowest circulating folate levels. Maternal plasma folate levels were positively associated with fetal weight (r = 0.712, p<0.01, Fig.1b). The mean daily diet and vitamins consumption can be found in Table 1. Based on diet composition, HF fed females had the lowest folate consumption.

**Fig. 1.**
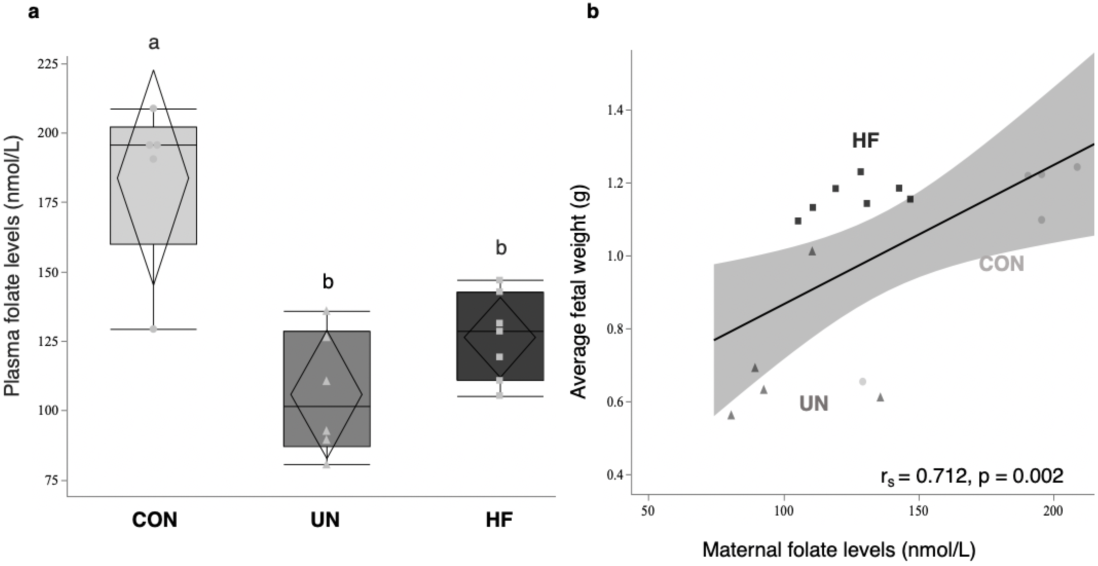
Plasma folate levels in CON, UN and HF mothers and association with fetal weight at GD18.5. Maternal plasma folate levels were lower in UN and HF mothers compared to CON (p<0.0001) (a). Data are quantile box plots with 95% confidence diamonds. Groups with different letters are significantly different (Tukey’s *post hoc*). Mean fetal weight was positively associated with maternal plasma folate levels (b). Data are linear regression with 95% confidence intervals (p<0.01). r_s_ = Spearman correlation coefficient. (Circle = CON, Triangle = UN, Square = HF)

### The maternal gut harbours an array of lactobacilli that are altered by HF diet

To determine the impact of maternal malnutrition on gut lactobacilli we measured the relative abundance of lactobacilli in maternal caecal contents. We first identified 15 maternal gut bacterial taxa (empirical operational taxonomic units (eOTU)) within the *Lactobacillus* genera in pregnant mice at GD18.5 (Fig. 2 a). Of these, HF fed mothers had a higher relative abundance of three specific lactobacilli eOTU compared to UN mothers (p<0.05, Fig. 2 c).

**Fig. 2.**
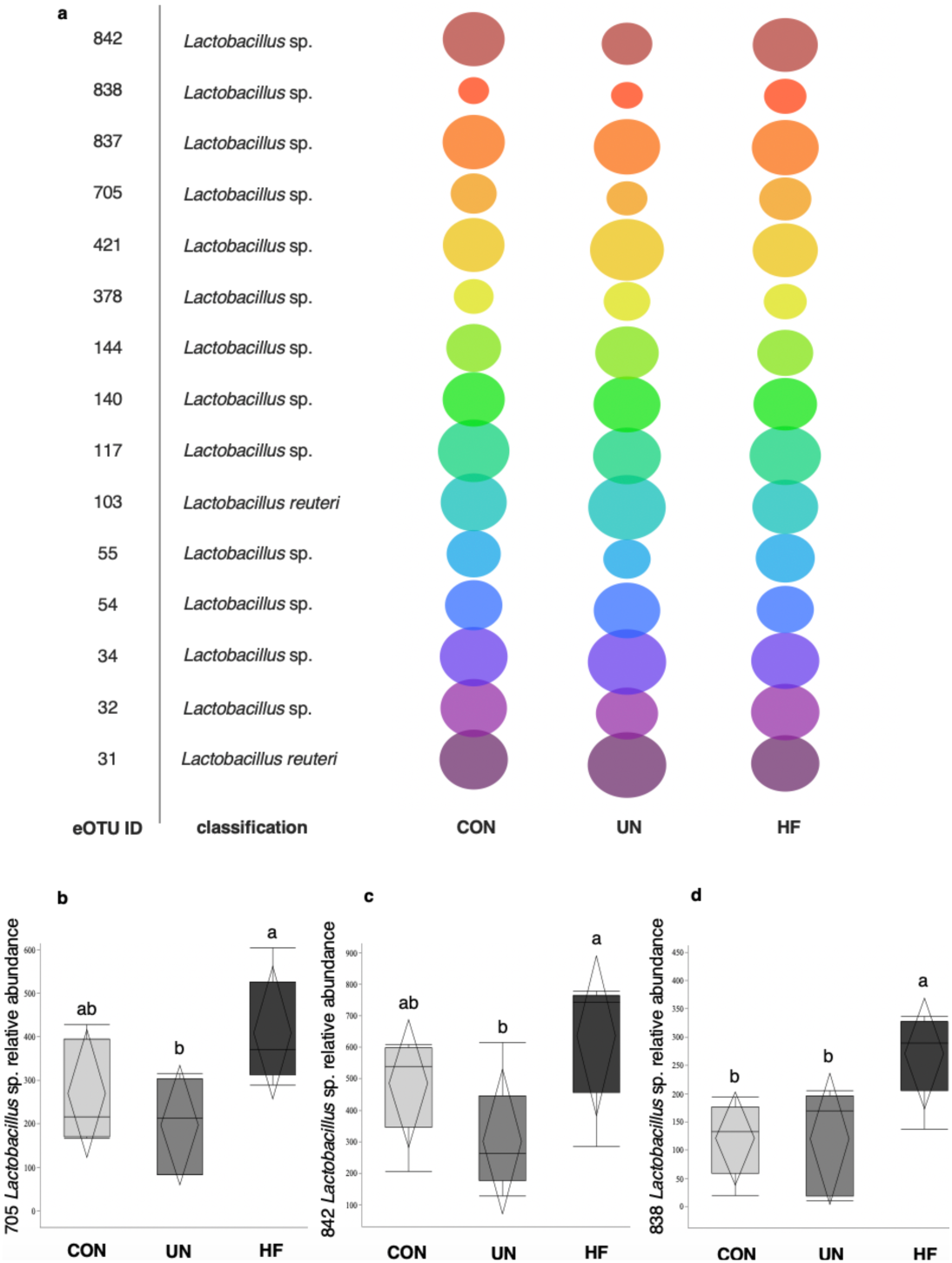
Array of lactobacilli in CON, UN, and HF maternal gut and abundance of lactobacilli in CON, UN and HF maternal gut. Bubble plot of relative abundance of 15 gut bacterial taxa (eOTU) within the *Lactobacillus* genus in pregnant mice at GD18.5 (a). Relative abundance of each eOTU is indicated by the size of the bubble. HF fed mothers had a higher relative abundance of three specific lactobacilli eOTU compared to mice fed an UN diet (a & b) (p<0.05) and compared to mice fed a CON and UN diet (c) (p<0.05) (b). Data are quantile box plots with 95% confidence diamonds. Groups with different letters are significantly different *(Tukey’s post hoc)*

### Maternal malnutrition is associated with altered expression of folate and inositol transporters in maternal SI

Maternal HF diet was associated with higher mRNA expression of the folate transporters *Pcft* compared to UN mothers (p<0.001, Fig. 3 a) and *Rfc1* compared to UN and CON mothers (p<0.01, Fig. 3 b). Inositol transporter *Smit2* mRNA expression levels were lower in the SI of HF fed compared to UN mothers (p<0.05, Fig. 3 c). There was no difference in *TonEBP* mRNA expression levels between the diet groups (Fig. 3 d). At the protein level, the cellular localisation of ir-PCFT and ir-SMIT2 was similar in all mothers, inclusive of diet. ir-PCFT staining was localised to the SI muscularis mucosae, mucosa, and the simple columnar epithelium surrounding the villi (Fig. 4 a). Semi-quantification of ir-PCFT staining intensity did not differ between the diet groups (Table 2). ir-SMIT2 staining was localised to the muscularis mucosae and simple columnar epithelium (Fig. 4 b). There was no difference in ir-SMIT2 staining intensity between diet groups (Table 2).

**Table 2.** PCFT and SMIT2 maternal SI staining intensity.

|  | CON | UN | HF | p-value |
| --- | --- | --- | --- | --- |
| **ir-PCFT in maternal SI** |  |  |  |  |
| Muscularis mucosae | 2.0 ± 1.45 | 2.1 ± 1.08 | 2.0 ± 1.45 | NS |
| Mucosa | 3.9 ± 0.10 | 2.4 ± 1.26 | 1.9 ± 1.85 | NS |
| Villi | 3.8 ± 0.25 | 2.6 ± 0.69 | 2.0 ± 1.46 | NS |
| **ir-SMIT2 in maternal SI** |  |  |  |  |
| Muscularis mucosae | 2.1 ± 1.71 | 1.9 ± 1.84 | 1.6 ± 1.54 | NS |
| Mucosa | 1.1 ± 1.92 | 1.3 ± 2.31 | 1.3 ± 1.63 | NS |
| Villi | 1.3 ± 2.31 | 1.7 ± 2.08 | 1.1 ± 1.20 | NS |

Data are mean (± standard deviation) intensity of ir-PCFT and ir-SMIT2. staining for the specified diet group determined from the average staining intensity across all six images for each SI tissue layer and each dam. Intensity was score at 0 = absent, 1 = weak, 2 = moderate, 3 = strong, 4 = very strong. Difference between diet groups determined at 5% level of significance. NS signifies no statistical difference between CON (n= 3), UN (n=3) and HF (n=5) within the specified tissue layer for both ir-PCFT and ir-SMIT2.

**Fig. 3.**
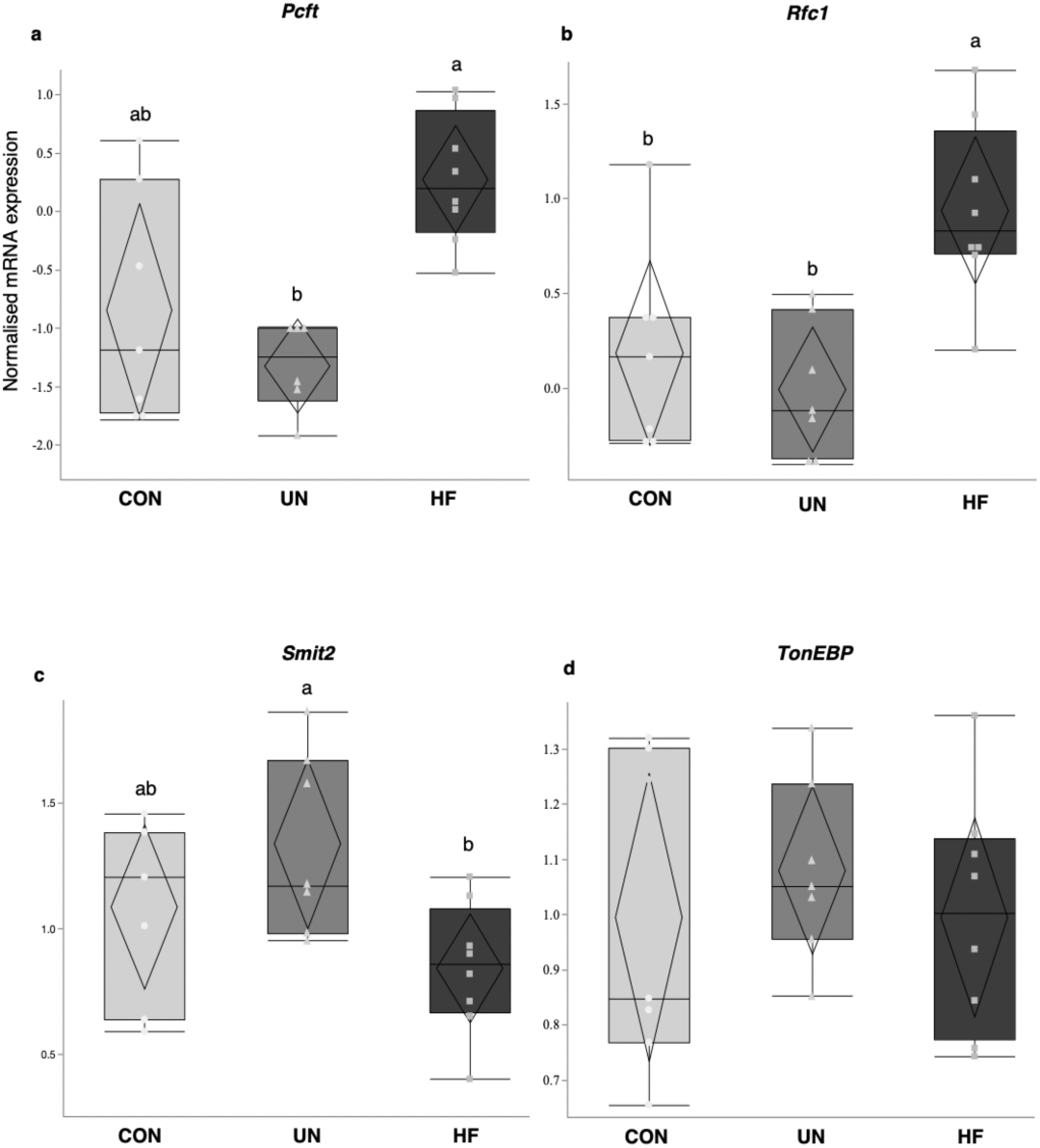
Maternal HF diet alters folate and inositol gene expression in the maternal SI. HF fed diet was associated with higher mRNA expression levels of the folate transporters *Pcft* compared to UN diet (p<0.001) (a) and *Rfc1* compared to UN and CON diet (p<0.01) (b). HF diet was associated with lower expression levels of inositol transporter *Smit2* compared to UN mothers (p<0.05) (c). There was no difference in expression levels of *TonEBP* between diet groups (d). Data are quantile box plots with 95% confidence diamonds. Groups with different letters are significantly different (Welch/Games-Howell or Tukey’s *post hoc*). (Circle = CON, Triangle = UN, Square = HF)

**Fig. 4.**
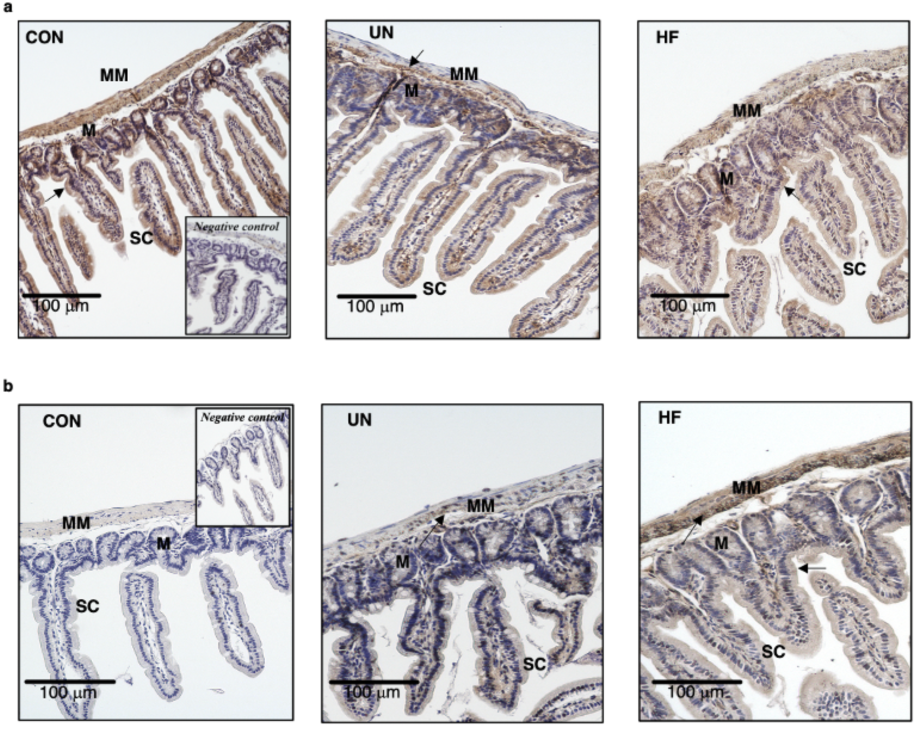
Representative images of immunoreactive PCFT (a) and SMIT2 (b) distribution in CON, UN and HF maternal SI samples with negative controls (inset). Images captured at 20X magnification, scale bar = 100 µm. ir-PCFT and SMIT2 distribution was not significantly different among CON (n=3/antibody), UN (n=3/antibody) and HF (n=5/antibody) maternal SI samples. Staining was localised to the muscularis mucosae (MM), mucosa (M), and simple columnar epithelium (SC) for ir-PCFT (a) and the muscularis mucosae (MM) and simple columnar epithelium (SC) for ir-SMIT2 (b). Arrows indicate areas where ir-PCFT and ir-SMIT2 staining is localized

### Maternal undernutrition is associated with decreased placental Frβ gene expression

Maternal UN was associated with reduced placental mRNA expression of *Fr*β, but not *Frα*, compared to CON and HF placentae (p<0.05, Fig. 5 d). There were no differences in mRNA expression levels of placental *Pcft, Rfc1, Frα, TonEBP*, and *Smit1* between the diet groups (Fig. 5). Fetal weight was negatively associated with placental *Fr*α mRNA expression (r=0.337, p<0.05, Fig. 6 a) and positively associated with *Fr*β mRNA expression (r=0.596, p<0.001, Fig. 6 b). When stratifying by sex, we found that HF male placentae had lower *Rfc1* mRNA expression compared to CON and that HF and CON male placentae had higher *Fr*β mRNA expression compared to UN (Online Resource Fig. 1 b & d). Female placenta had higher *Fr*β mRNA expression compared to CON and UN (Online Resource Fig. 1 d). At the protein level, the cellular localisation of ir-FRβ was similar in all placentae, inclusive of diet. ir-FRβ staining was localized to the decidua, junctional and labyrinth zones, with most intense staining appearing in the labyrinth zone (Fig. 7), and staining intensity did not differ between the diet groups (Table 3).

**Table 3.** FRβ placenta staining intensity.

|  | CON | UN | HF | p-value |
| --- | --- | --- | --- | --- |
| **ir-*FRβ* in placenta** |  |  |  |  |
| Decidua | 2.1 ± 1.35 | 3.0 ± 1.00 | 3.0 ± 1.04 | NS |
| Junctional zone | 2.1 ± 1.30 | 3.2 ± 0.96 | 3.2 ± 0.92 | NS |
| Labyrinth zone | 1.8 ± 1.68 | 2.4 ± 1.39 | 3.1 ± 1.03 | NS |
| Chronic plate | 1.9 ± 1.56 | 2.3 ± 1.59 | 2.5 ± 1.40 | NS |

Data are mean ( ± standard deviation) intensity of ir-*FRβ* staining for the specified diet group determined from the average staining intensity across all six images for each placental tissue layer and each animal. Intensity was score at 0 = absent, 1 = weak, 2 = moderate, 3 = strong, 4 = very strong. Difference between diet groups determined at 5% level of significance. NS signifies no statistical difference between CON (n=8), UN (n=8) and HF (n=8) within the specified tissue layer.

**Fig. 5.**
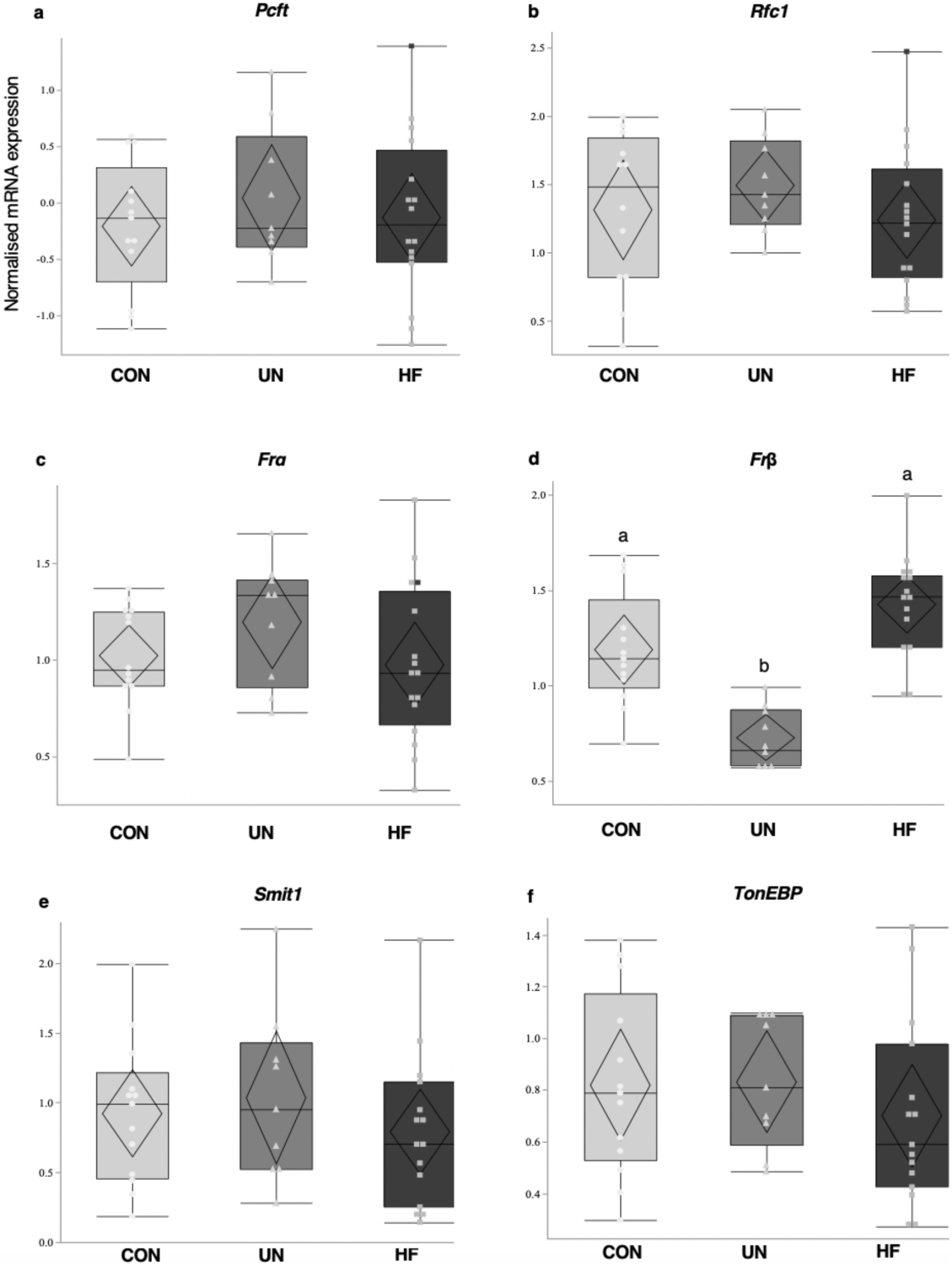
Folate and inositol transporter mRNA expression in CON, UN and HF placentae. *Fr*β mRNA expression was reduced in UN placentae compared to CON and HF placentae (p<0.05) (d). Data are quantile box plots with 95% confidence diamonds. Groups with different letters are significantly different (Tukey’s *post hoc*). (Circle = CON, Triangle = UN, Square = HF)

**Fig. 6.**
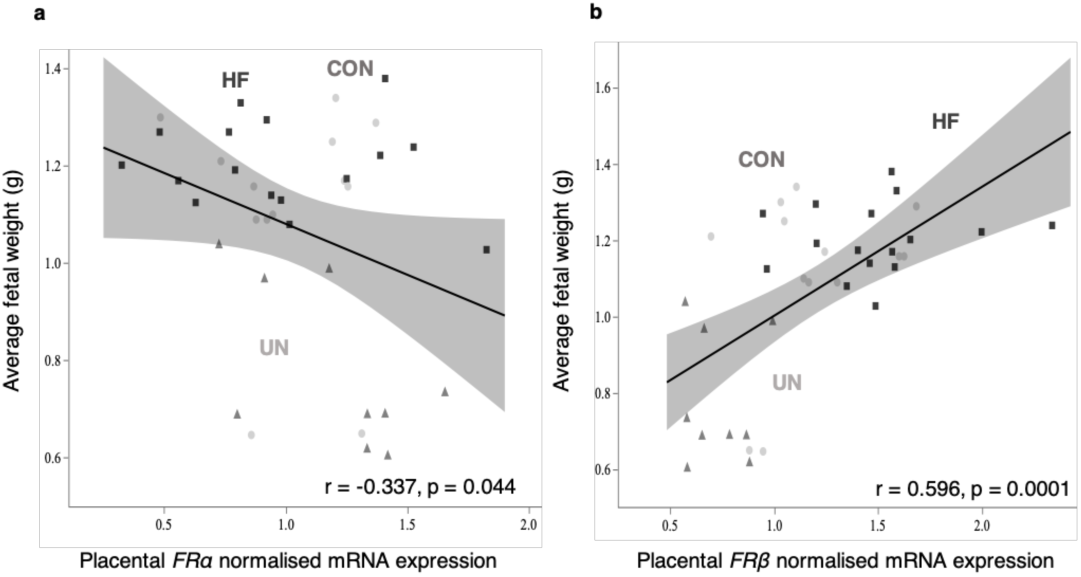
Association between fetal weight and folate receptor *α* and β mRNA expression at GD18.5. Fetal weight was negatively associated with placental *Frα* expression (a) and positively associated with *Fr*β expression (b). Data are linear regression with 95% confidence intervals, p<0.05. r = Pearson correlation coefficient. (Circle = CON, Triangle = UN, Square = HF)

**Fig. 7.**
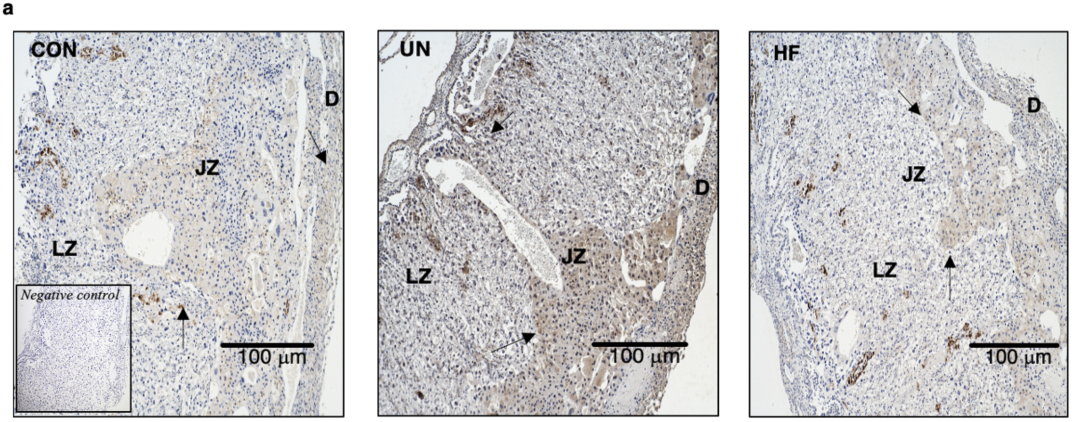
Representative images of immunoreactive FRβ distribution in CON, UN and HF placentae with negative controls (inset). Images captured at 20X magnification, scale bar = 100 µm. ir-FRβ distribution was not significantly different among CON, UN and HF placentae. Staining was localized to the decidua (D), junctional (JZ), and labyrinth zones (LZ). Arrows indicate areas where ir-FRβ staining is localized

### Maternal malnutrition is associated with altered folate and inositol gene expression in the fetal gut

Consistent with gene expression changes in the maternal gut, fetuses from HF fed mothers had higher gut mRNA levels of the folate transporter *Pcft* compared to fetuses from CON and UN mothers (p<0.0001, Fig. 8 a) and higher *Frα* compared to fetuses from UN mothers (p<0.01, Fig. 8 b). In contrast to the maternal SI, fetuses from HF fed mothers had higher gut inositol transporter *Smit2* mRNA expression compared to fetuses from CON and UN mothers (p<0.01, Fig. 8 c). There were no differences in mRNA expression levels of fetal gut *TonEBP* between the diet groups (Fig. 8 d). When stratifying by sex, we found that both HF female and male fetuses had higher gut mRNA expression of *Pcft* compared to CON and UN (Online Resource Fig. 2 a). *Frα* mRNA expression was higher in HF male fetal gut compared to CON and UN male fetal gut (Online Resource Fig. 2 b) and *Smit2* mRNA expression was higher in HF female fetal gut compared to CON female fetal gut (Online Resource Fig. 2c).

**Fig. 8.**
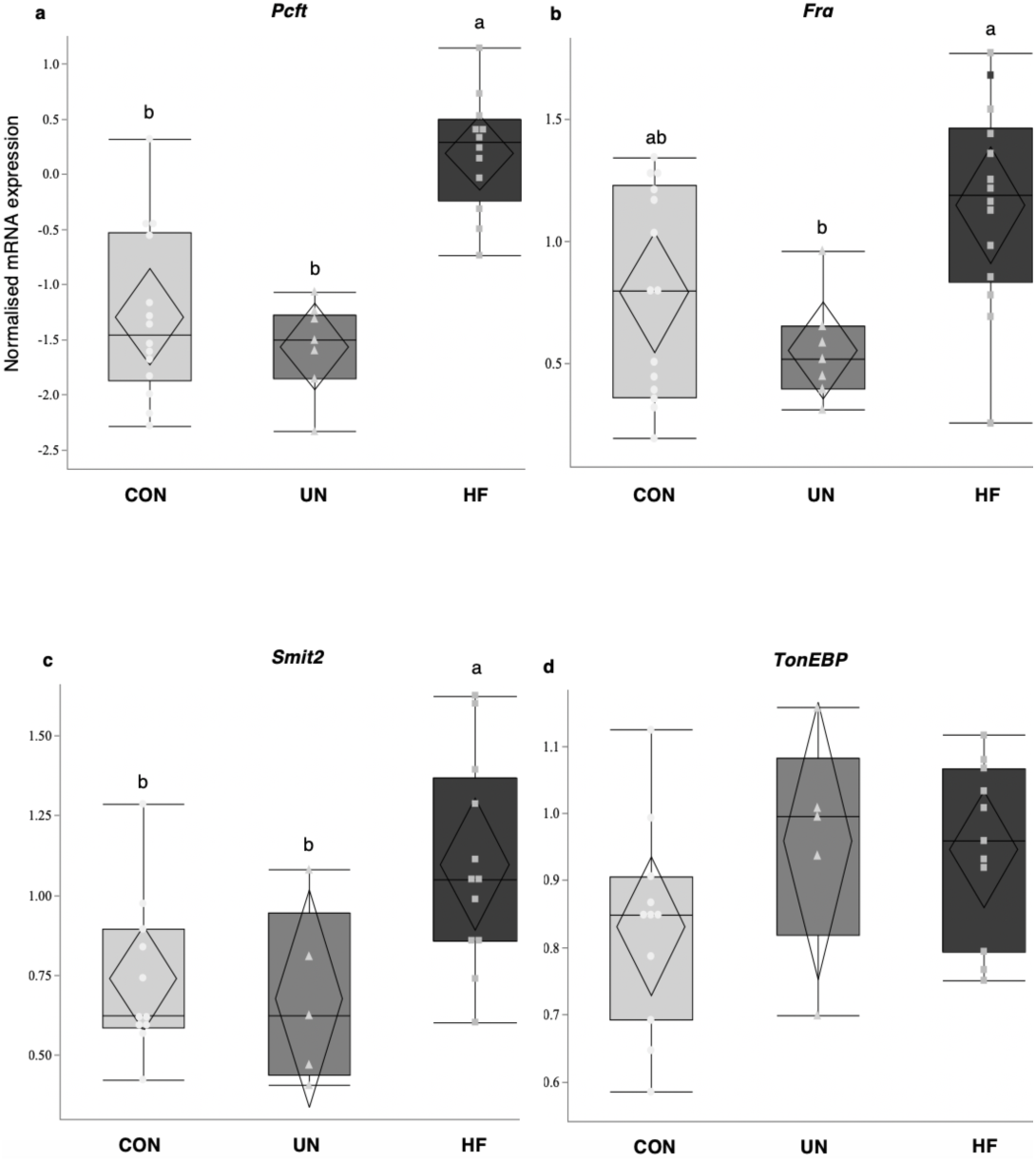
Maternal malnutrition alters folate/inositol gene expression in the fetal gut. Fetuses from HF fed mothers had higher mRNA expression levels of the folate transporters *Pcft* compared to CON and UN (p<0.0001) (a), and higher *Frα* compared to UN (p<0.01) (b). Fetuses from HF fed mothers had higher expression levels of the inositol transporter *Smit2* compared to UN (p<0.01) (c). There was no difference in expression levels of the inositol transporter *TonEBP* between diet groups (d). Data are quantile box plots with 95% confidence diamonds. Groups with different letters are significantly different (Tukey’s *post hoc*). (Circle = CON, Triangle = UN, Square = HF)

### Maternal diet coordinates micronutrient pathways across the nutritional pipeline

To visualise our maternal and fetal outcome measures and determine whether, and which, specific features could discriminate mothers from each other, we performed principal component analysis (PCA). Overall, PCAs revealed that clustering was largely determined by maternal nutritional exposure. Our first PCA included the maternal features plasma folate levels, relative abundance of significant maternal gut lactobacilli (eOTU) and mRNA levels of folate and inositol genes. Distinct clusters of mothers emerged primarily based on PC1, where HF mothers clustered separately from CON and UN (Fig. 9 a). The features considered important for PC1 were represented by *Pcft* and *Rfc1* mRNA expression and the three significantly different lactobacilli (eigenvalue [EV] 3.10, explaining 44.3% of the variance in the data; Online Resource Fig. 3 a). Based on PC2, the features with the greatest EVs were represented by *Pcft* and *Smit2* mRNA expression (EV 1.35, explaining 19.3% of the variance in the data; Online Resource Fig. 3 a). Our second PCA included the fetal features placental and gut folate/inositol mRNA levels and fetal and placental weights. Fetuses from HF mothers clustered separately from UN, whereas fetuses from CON mothers were intermingled; clustering was primarily based on PC1 (Fig. 9 b). The features considered important for PC1 were represented by fetal weight, *Pcft, Frα*, and *Smit2* gut mRNA expression and *Fr*β placental mRNA expression (EV 2.74, explaining 45.9% of the variance in the data) and for PC2 were represented by *Pcft, Smit2* gut mRNA expression and *Fr*β placental mRNA expression (EV <1, explaining 19.8% of the variation; Online Resource Fig. 3 b).

**Fig. 9.**
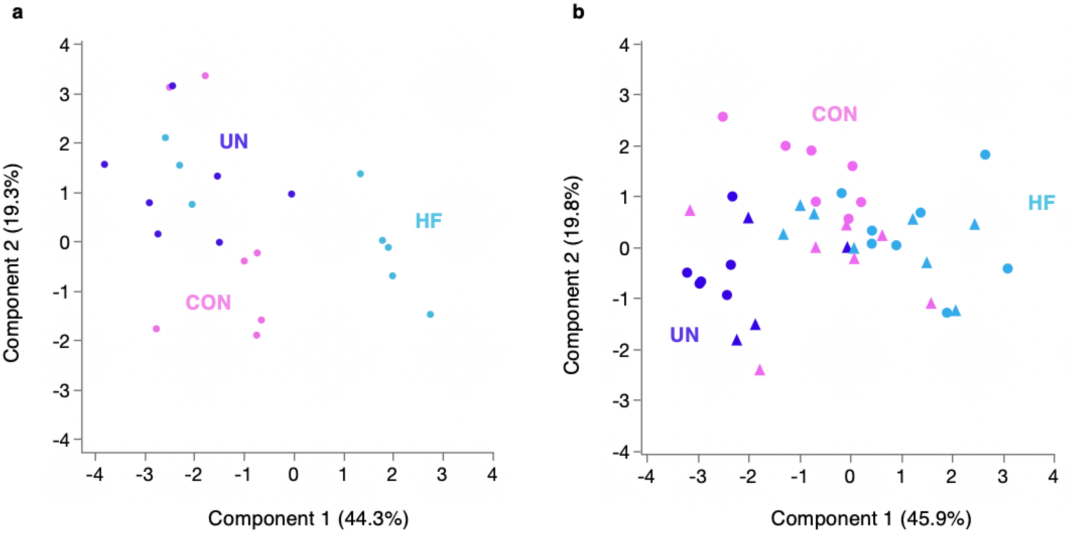
Principal component analysis (PCA) of maternal (a) and fetal (b) variables. HF fed mothers are separated from CON and UN mothers. The first component explains 44.3% of the variation and the second component 19.3% (a). Fetuses from HF fed mothers are separated from UN; fetuses from CON are mixed. The first component explains 45.9% of the variation and the second component 19.8% (b). (Triangle = Female, Circle = Male)

## Discussion

We used a systems-physiology approach, considering the maternal holobiont (the host and its microbiota), placenta and fetus, to evaluate the effects of diet-induced changes in pregnancy on micronutrient pathways that are important for shaping fetal growth and development. As a first step, we sought to answer three questions: does maternal malnutrition alter microbes in the maternal gut that could be important for folate production? Does malnutrition impact maternal micronutrient levels and folate and inositol transport in the maternal gut and placenta? And does malnutrition affect gut folate and inositol pathways in the developing fetus, which may contribute to the programming of gut dysfunction long-term? We found that both types of malnutrition resulted in lower circulating folate levels, but they affected folate and inositol pathways across the nutritional pipeline differently: gut lactobacilli relative abundance levels and maternal gut expression of *Pcft* and *Rfc1* were higher, but *Smit2* lower, in HF mothers compared to UN and CON. Among fetuses, placental expression of *Fr*β was higher in HF placentae compared to UN and gut expression of *Frα* and *Smit2* was higher in fetuses from HF fed mothers compared to UN and CON.

In addition to placental function, an important aspect of our study was to interrogate maternal gut function. Our finding that maternal SI *Pcft* and *Rfc1*expression is responsive to dietary intakes, even at nutrient intake levels that are considered adequate or better, is consistent with reports that low folate intake is associated with upregulation of folate transporters [53]. Further, we found that HF fed mothers had a higher relative abundance of three specific eOTU within the *Lactobacillus* genus compared to UN mothers, results that are in part consistent with our hypothesis that malnutrition would alter levels of maternal gut lactobacilli. Despite these findings, both HF and UN mothers had lower circulating folate levels, consistent with our observation that UN and HF fed mice consumed less, albeit still adequate, folate from their diets, and consistent with reports of lower circulating folate levels with maternal underweight and obesity [7–11]. In the current study, we do not have direct evidence for reduced maternal gut bacterial production of folate impacting host micronutrient levels, and thus, the specific links between host folate status and gut bacterial folate production have yet to be disentangled. Whilst *Lactobacillus* is a dominant genus of microbes in the mouse gut, we could not detect any species in the *Bifidobacterium* genus [54]. Species in the latter genus also contribute to folate production in the host and are a major component of the mouse microbiome [32,34]. The influence of folate-producing microbes on the expression and activity of receptors and transporters like *Rfc1* and *Pcft*, that play dominant roles in intestinal folate absorption and uptake/transport respectively, will be a key concept to explore in the context of pregnancy and folate-mediated fetal growth restriction and disorders [16,55].

Further, our study highlights an important relationship between maternal malnutrition and placental folate/inositol transporters. We found that UN placentae had lower *Fr*β mRNA expression levels compared to CON and HF placentae, whilst there was no difference in the expression of the major placental folate receptors and transporters *Frα, Pcft*, and *Rfc1* between the diet groups. Additionally, although we report lower *Smit2* mRNA expression in maternal SI of HF mothers, we did not observe changes in placental expression levels of inositol transporters. As we were not able to measure circulating inositol levels in mothers and levels in the diets were not defined by manufacturers, it may be that, as with folate content, inositol content varied between the diets. Based on these findings, we cannot speculate about the cause of reduced SI *Smit2* or its physiologic relevance, or whether the absence of a dietary effect on placental inositol transport means this micronutrient is of lesser importance to overall fetal development at the end of pregnancy. Nevertheless, our data suggest that the placenta sufficiently buffers the suboptimal pregnancy environments established by UN and HF exposures, at least with respect to folate and inositol uptake and transport, likely in an effort to maintain fetal growth and development under these conditions [46,48]. As placental expression of folate transporters is dynamic across pregnancy, future studies should examine the ontogeny of B vitamin transporter expression to fully understand the critical periods during which diet-induced changes in pregnancy phenotypes can alter folate and inositol transport [18,19].

Importantly, to our knowledge, this is the first study to measure mRNA levels of folate and inositol transporters in the fetal gut and show early life programming of offspring gut micronutrient pathways by exposure to malnutrition *in utero*. Fetuses from HF fed mothers had higher gut mRNA expression levels of folate transporters *Pcft* and *Frα* and inositol transporter *Smit2*. Whilst it has yet to be conclusively determined if bacterial colonisation of the fetal gut occurs in utero, species in the *Lactobacillus* genus are amongst the first microbes to colonise the infant gut after vaginal birth [56].Therefore, the presence of folate and inositol transporters in the fetal gut may be a preparatory step to complement the micronutrient production that is likely to occur after bacterial colonisation in the early newborn gut. However, dietary reprogramming of these key transporters *in utero*, may have both short- and long-term implications for their function and ultimately, host micronutrient status. This will be a critical concept to explore, including whether these findings are more broadly observed in other gut nutrient transport systems in fetal and neonatal life, and if they are causal of poor growth trajectories after birth.

A limitation to our study is its cross-sectional design. It remains to be determined whether maternal malnutrition impacts gut bacterial folate production and intestinal and placental folate and inositol gene/protein expression in very early pregnancy. That said, our findings are still relevant since suboptimal micronutrient status, production and transport of folate and inositol at the end of pregnancy could be a marker of *in utero* events earlier in pregnancy, and a window to future maternal and offspring health. Additionally, the CON/UN and HF diets used had differences in their nutritional composition (neither diet indicated the amount of inositol). While not the purpose of our study, future studies should explore how differences in micro/ macronutrient levels impact the maternal holobiont, placenta, and fetus, with respect to micronutrient production and transport pathways, by controlling for specific micronutrients. Here, our aim was to use established diets that elicit specific pregnancy phenotypes to examine the subsequent effect on circulating folate levels and nutrient transport physiology across the gut-placental-fetal pipeline. For example, while our HF model had a high proportion of calories from fat, it did not result in fetal growth restriction as seen in many other animal models, which is not entirely reflective of the human situation [57]. That said, our HF model does parallel diets from populations who do consume higher amounts of fat, such as the Alaskan Inland Inuit populations who have undergone nutritional adaptations (such as high carbohydrate, low protein diets that are regularly over 40% fat under normal conditions) due to less than favourable environmental changes, and populations residing in the Pacific Islands, who have high levels of daily absolute fat intake and saturated fat intake as a result of various social determinants (e.g. availability of processed foods and access to fresh local food) [58,59]. Furthermore, maternal plasma inositol concentrations could not be measured due to limited sample volume, which was prioritised for measuring folate. Similarly, fetal plasma folate concentrations could not be measured due to limited sample volume, therefore we do not know the effects of these adverse exposures on circulating micronutrient levels in the fetus at term. However, others have shown that maternal diets deficient in folate, resulting in low maternal serum folate levels, impact fetal growth at the end of gestation (GD 18.5) [60].

Our systems physiology approach is an important advance to better understand the (micro)nutrient pathways that regulate fetal growth and development, and contrasts with other studies that only consider the placenta[1,2]. Our findings provide a first step to more complete insight into microbial and micronutrient physiology in pregnancy and, once carefully characterised across pregnancy and if translated to human cohorts, may aid in efforts to identify new biomarkers and therapeutic targets for women and pregnancies at high-risk of micronutrient-associated fetal disorders or poor fetal growth and development.

## Acknowledgments

We would like to thank Amanda J. MacFarlane from the Nutrition Research Division at Health Canada for her contributions to this manuscript including to the methods, investigation, data interpretation and manuscript revisions and Nathalie Behan for completing the microbiological folate assays. We deeply thank Professor Stephen Lye for resource contributions towards the animal model. This research is funded by the Faculty of Science, Carleton University and TVM is the recipient of a merit award from the department of Obstetrics and Gynaecology at Mount Sinai Hospital.

## Author Contributions and Notes

Conceptualisation: Kristin L Connor, Elia Palladino; Methodology: Kristin L Connor, Elia Palladino; Formal analysis and investigation: Elia Palladino, Kristin L Connor; Writing - original draft preparation: Elia Palladino, Kristin L Connor; Writing - review and editing: Tim Van Mieghem, Kristin L Connor, Elia Palladino. Funding acquisition: Kristin L Connor; Resources: Kristin L Connor; Supervision: Kristin L Connor.

---

**Online Resource Table 1.**
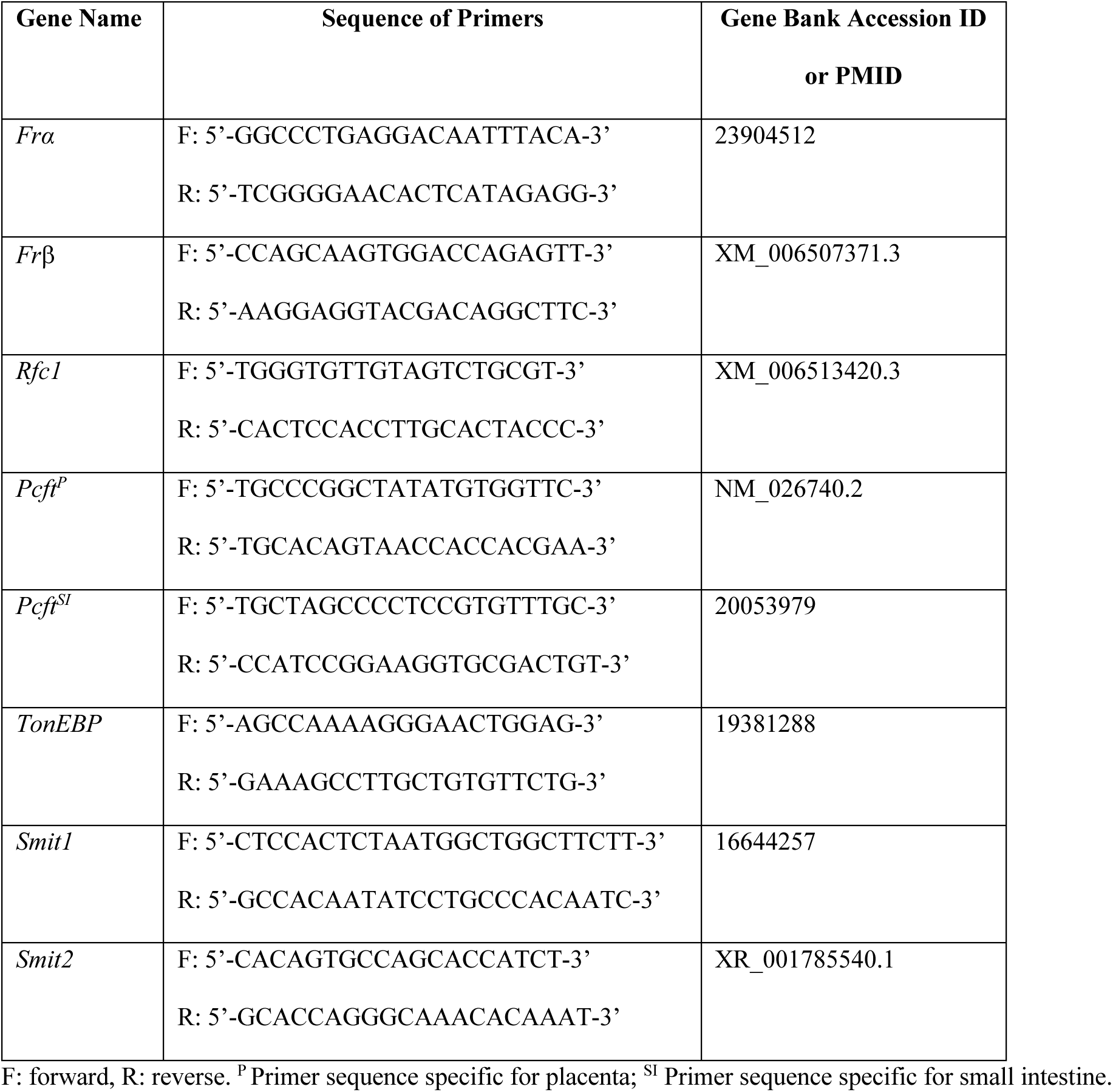
Primer sequences for quantitative PCR (qPCR).

**Online Resource Fig. 1.**
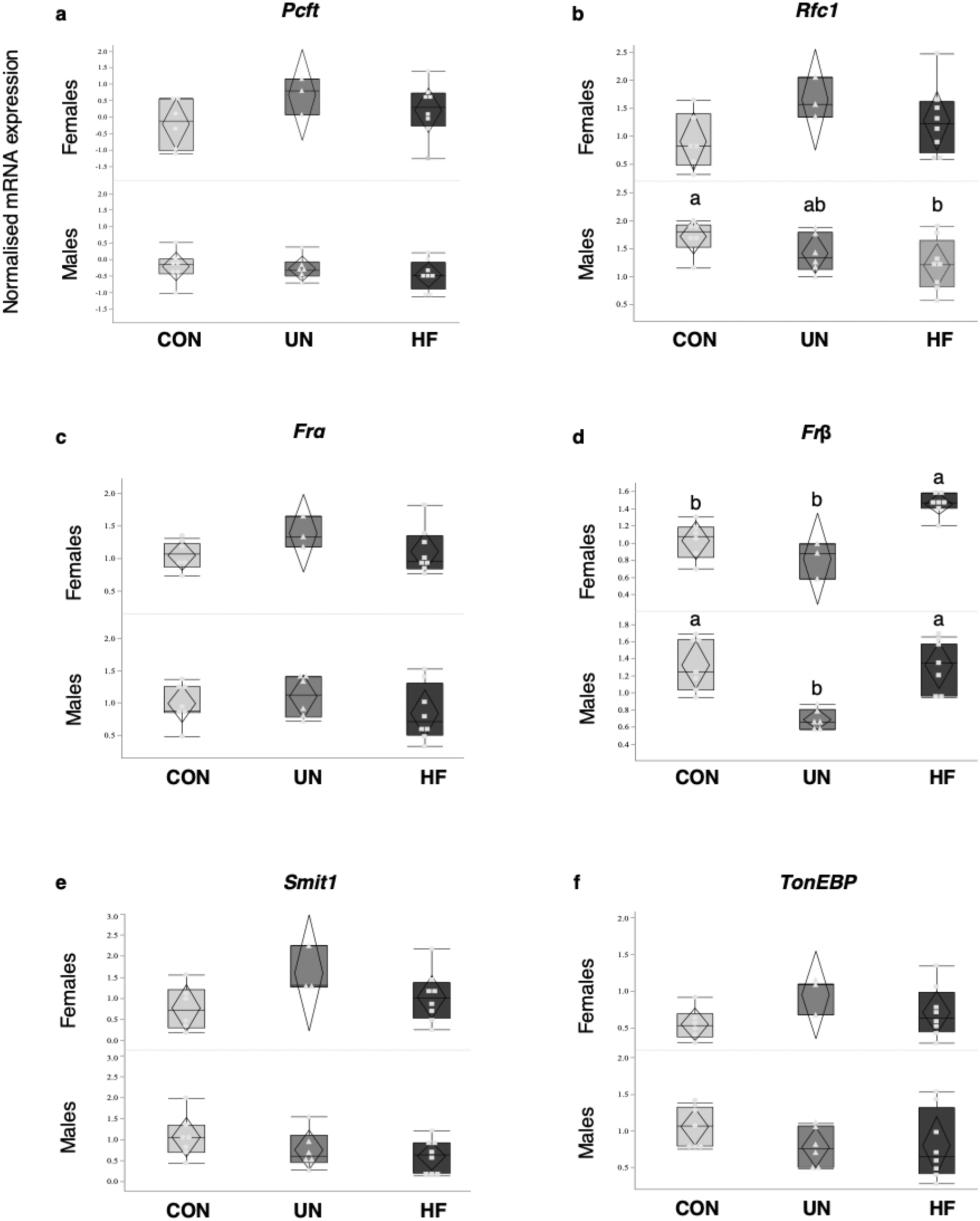
Folate and Inositol transporter mRNA expression in CON, UN. and HF placentae stratified by fetal sex *Rfcl* mRNA expression was reduced in HF male placentae compared to CON male placentae (p<0.0755) (B). *Frβ* mRNA expression was higher in HF female placentae compared to CON and UN female placentae (p<0.001) and higher in CON and HF male placentae compared to UN (p<0.00l) (D). Data are quantile Cox pots with 95% confidence diamonds. Groups with different letters are significantly different (Tukey’s *post hoc*). (Circle = CON, Triangle = UN, Square = HF).

**Online Resource Fig. 2.**
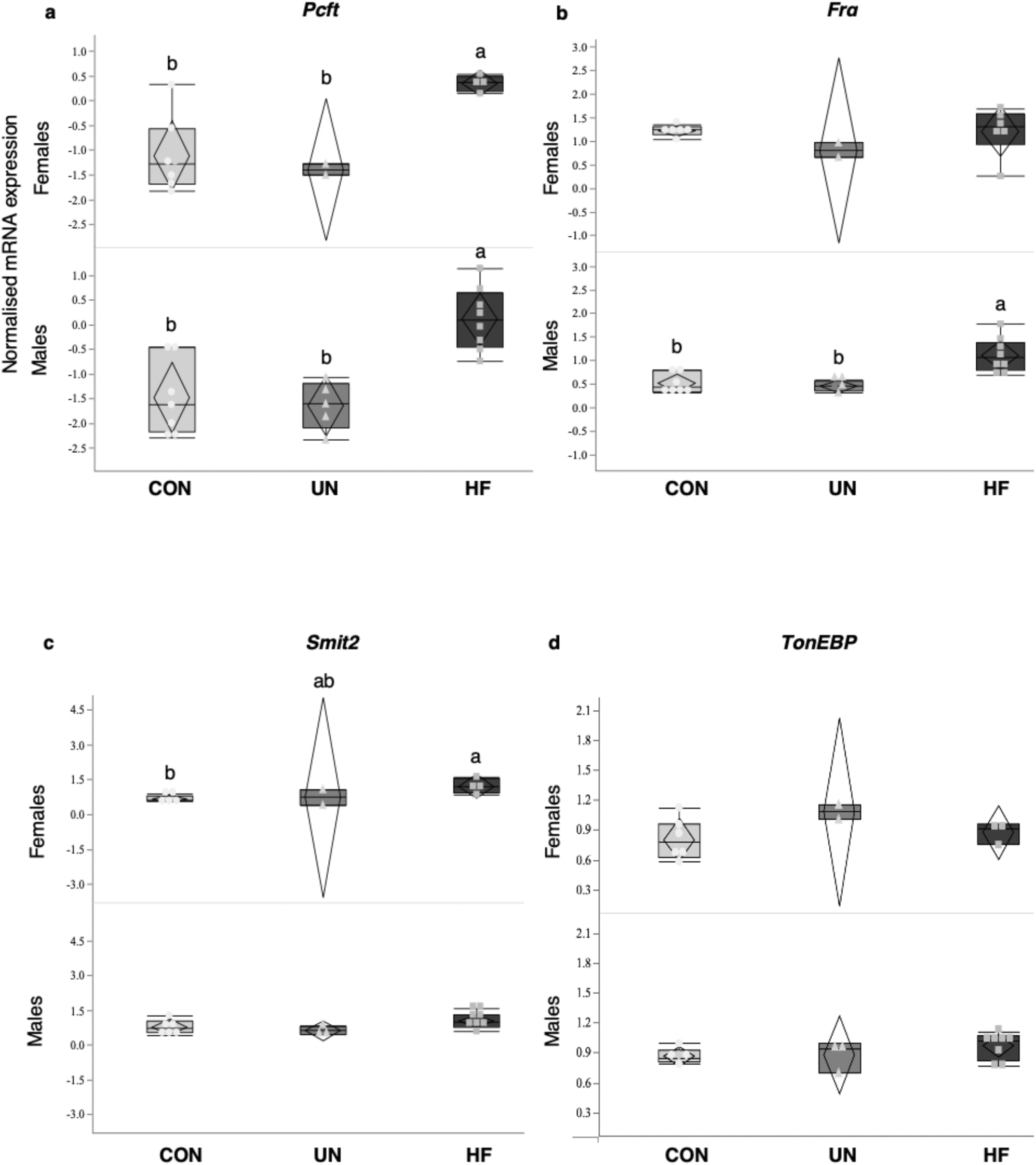
Folate and Inositol transporter mRNA expression in CON, UN, and HF fetal gut stratified by fetal sex. *Pcft* mRNA expression was higher in HF female and male fetal gut compared to CON and UN fetal gut (p<0.05 for female; p<0.001 for male) (A). *Frα* mRNA expression was higher in HF male fetal gut compared to CON and UN male fetal gut (p<0.00l) (B). *Smrt2* mRNA expression was higher in HF female fetal gut compared to CON female fetal gut (p<0.05) (C). Data are quantile box plots with 95% confidence diamonds Groups with different letters are significantly different (Tukey’s *post hoc*). (Circle = CON, Triangle = UN, Square = HF).

**Online Resource Fig. 3.**
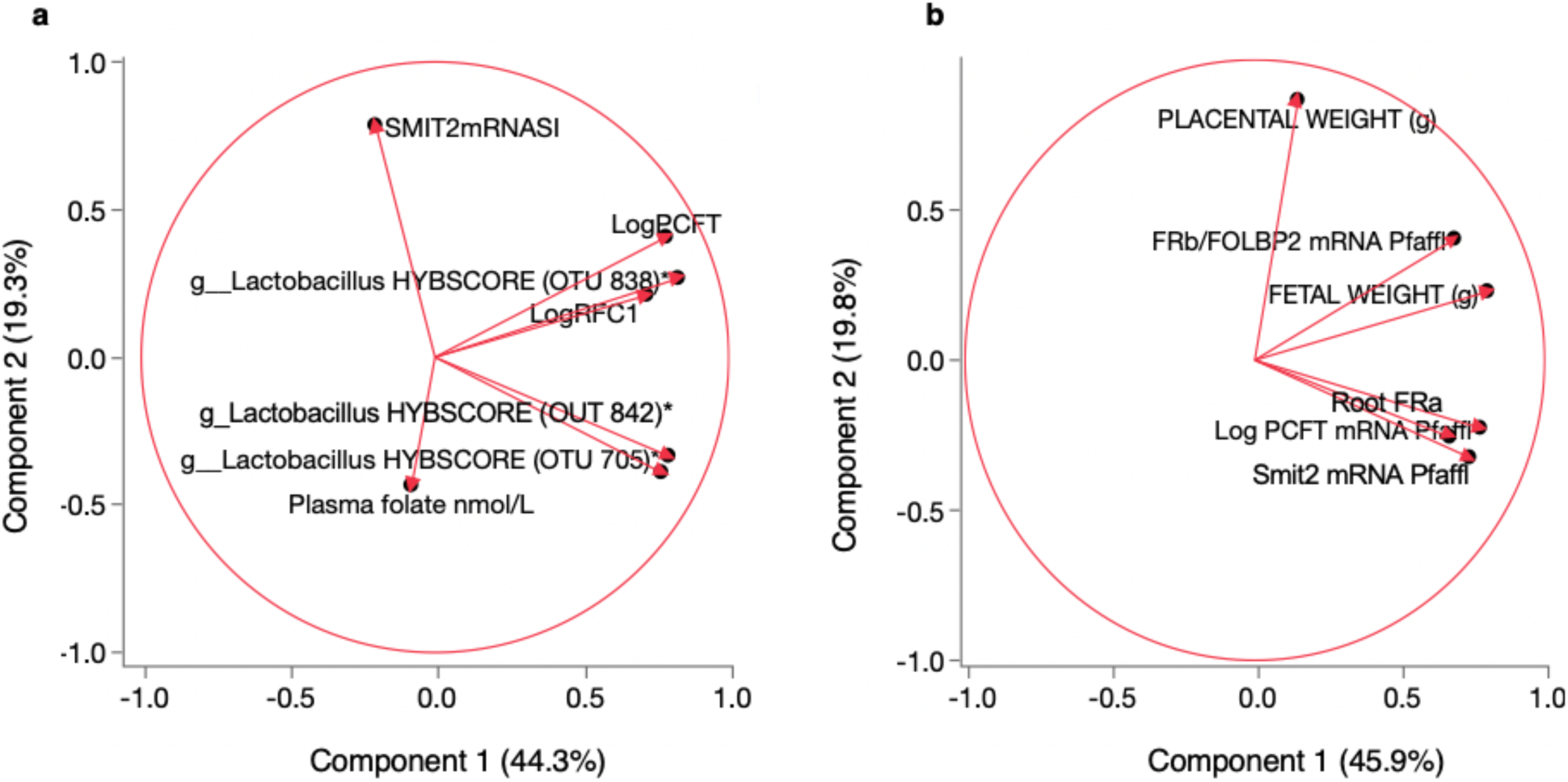
Correlations determined by PCA of maternal (A) and fetal (B) variables. (A) *Pcft* anti *Rfc1* mRNA expression and eOTU 838 are correlated. Maternal plasma folate levels and *Smit2* mRNA expression were negatively correlated. (B) *Pcft. Frα. Smit2* mRNA expression were correlated and *Frβ* and fetal weight (g) were positively correlated.

